# Endogenous retroviruses activate MARCO-mediated inflammatory response to block retroviral infection

**DOI:** 10.1101/2024.09.03.610969

**Authors:** Xuming Hu, Wang Guo, Huixian Wu, Jinlu Liu, Xujing Chen, Xiao Han, Yu Zhang, Yang Zhang, Zhengfeng Cao, Qiang Bao, Wenxian Chai, Shihao Chen, Wenming Zhao, Guohong Chen, Hengmi Cui, Qi Xu

## Abstract

Endogenous retroviruses (ERVs) are remnants of ancient retroviral infections and can profoundly affect the host antiviral innate immune response, although the mechanisms by which these changes occur are largely unknown. Here we report that chicken-specific ERVs exert genetic resistance to exogenous retrovirus infection. Mechanistically, chicken-specific ERVs activated the scavenger receptor MARCO (macrophage receptor with collagenous structure)-mediated TLR3-IL-1β inflammatory response in macrophages. Under the presence of MARCO, macrophages response to viral infection through inducing TLR3-IL-1β inflammatory response. Conversely, lack of MARCO increased the viral replication levels and attenuated the antiviral inflammatory response. MARCO-mediated ligand delivery enhances TLR3-IL-1β antiviral response, and IL-1β expression is responsible for viral inhibition. Restoring MARCO or IL-1β expression overcomes viral infection in macrophages. Our study provides new insights into the molecular mechanisms underlying the host defense against exogenous retroviruses infection and may have important implications for the development of novel therapeutic strategies against retroviruses infection.

## Introduction

Endogenous retroviruses (ERVs) are remnants of ancient retroviral infections that have integrated into the host genome and are extensively prevalent in mammalian and chicken genomes (Stoye, 2012). Although initially considered “junk DNA”, it has recently attracted enormous attention owing to their important physiological functions in genome evolution (Biemont & Vieira, 2006; Cordaux & Batzer, 2009), embryonic development (Grow *et al*, 2015), tissue homeostasis and aging (Lima-Junior *et al*, 2021; Liu *et al*, 2023). Accumulated evidence has also highlighted their potential involvement in viral infection and immune modulation.

Mounting evidences support the idea that ERVs promote interferon production through several key protein sensors, including retinoic acid-inducible gene I (RIG-I) and melanoma differentiation-associated protein 5 (MDA5) (Chiappinelli *et al*, 2015; Licht, 2015; Roulois *et al*, 2015). These findings also raise the intriguing possibility that ERVs protect human embryos against different classes of viruses sensitive to the interferon induced transmembrane protein 1 (IFITM1)-type restriction (Grow *et al*, 2015). It has been shown that ERVs have shaped the evolution of a transcriptional network underlying the interferon response (Chuong *et al*, 2016). Study further reveals that tumor innate immunity primed by specific interferon-stimulated ERVs (Canadas *et al*, 2018). Together, these observations implicate ERVs play a key role in the interferon response, a major branch of innate immunity.

ERVs play an important role not only in the interferon response, but also in modulating inflammatory response, another major branch of innate immunity. The innate immune system acts as the first line of defense against invading pathogens and plays a crucial role in maintaining homeostasis. It has shown that ERVs promote tissue homeostasis and inflammation responses to skin microbiome through the cyclic GMP-AMP synthase (cGAS)/stimulator of interferon genes protein (STING) signalling (Lima-Junior *et al*, 2021). However, excessive viral mimicry generated by motivated ERVs promoted bowel inflammation through triggering the Z-DNA-binding protein 1 (ZBP1)-dependent necroptosis (Jiao *et al*, 2020; Wang *et al*, 2020). More importantly, the antiviral innate immunity functions of in the pro-inflammatory macrophages might be associated with ERVs-derived long noncoding RNAs (Luo *et al*, 2022; Zhou *et al*, 2019). These findings raising intriguing questions about ERVs contribution to immune homeostasis and disease pathogenesis through inflammatory response. However, the pro-inflammatory mechanisms of ERVs on immune cells, especially macrophages, are still not fully elucidated.

The above findings fully elucidate the role of ERVs themselves in antiviral innate immunity. ERVs have similar sequences to exogenous retroviruses, it is not clear how these ERVs sequences exert antiviral effects in response to exogenous retroviruses infection. Several studies believed that immune system may have co-opted its ERVs as a way to communicate with the exogenous microbiome (Hurst & Magiorkinis, 2015; Kassiotis & Stoye, 2016; Lima-Junior *et al*, 2021). However, the complex relationships about how exogenous retroviruses interfere with ERVs-mediated antiviral innate immunity remain largely unexplored. Currently, ERVs are broadly classified into classes I, II, and III based on the relatedness to exogenous retroviruses genera (Bolisetty *et al*, 2012; Jern *et al*, 2005). Class II chicken ERVs (chERVs) are the only ERVs that are highly homologous to the alpha retroviruses avian leukosis virus (ALV), which is the only known chicken retrovirus with both exogenous and endogenous activity (Bolisetty *et al*, 2012; Mason *et al*, 2020). This particularity suggesting chicken retroviral evolution may be different from that of other vertebrates. Therefore, chicken-specific ERVs provides a unique model for investigating the interaction between endogenous and exogenous retroviruses as well as the symbiotic relationship and interplay between ERVs and innate immunity.

Here we revealed an uncharacterized role of endogenous retroviruses in the inflammatory response and host genetic resistance using the unique model chicken-specific ERVs and CRISPR/Cas9 technology. In particular, chERVs can inhibit exogenous retroviral replication through activating the macrophage scavenger receptor MARCO-mediated inflammatory response. The present study reveals that chERVs and MARCO as genetic resistance factors and drug targets for developing antiviral strategies against exogenous retroviral infection.

## Results

### chERVs exert resistance to homologous exogenous ALV-J infection

To determine the host genetic resistance of chERVs to viral infection, we first investigated the expression pattern and antiviral function of chERVs in chickens infected by the avian leukosis virus subgroup J (ALV-J). We detected the expression level of chERVs-*env* transcripts in the central immune organs (spleen and bursa) of 8 chicken lines. The results showed that the expression level of chERVs-*env* transcripts in chicken line E immune organs (spleen and bursa) was abnormally higher than that of others chicken (Figure 1A-D). Next, we selected line B, E and H for antiviral experiments based on the distribution model of chERVs expression. Viral infection experiments demonstrated that chERVs exert genetic resistance to ALV-J infection. Compared to the chERVs-*env* transcripts-high chickens inoculated with ALV-J for 10 days, the chERVs-*env* transcripts-low chickens exhibited typical pathological changes. The most noticeable changes were observed in the yellow color of livers, enlargement of spleen and atrophy of bursa (Figure 1E-G). These changes are important clinical signs of ALV-J infection in chickens. Specifically, we also observed a significant reduction in the replication levels of ALV-J in the spleen and bursa of chERVs-*env*-high chickens compared to chERVs-*env*-low chickens (Figure 1H-K). These data suggest that chERVs-*env* transcripts play an important role in limiting the replication of ALV-J in chickens. In particular, chERVs-*env* transcripts may be used as a biomarker for ALV-J resistant chickens and potential targets for the development of new treatments for ALV-J infection.

**Figure 1.**
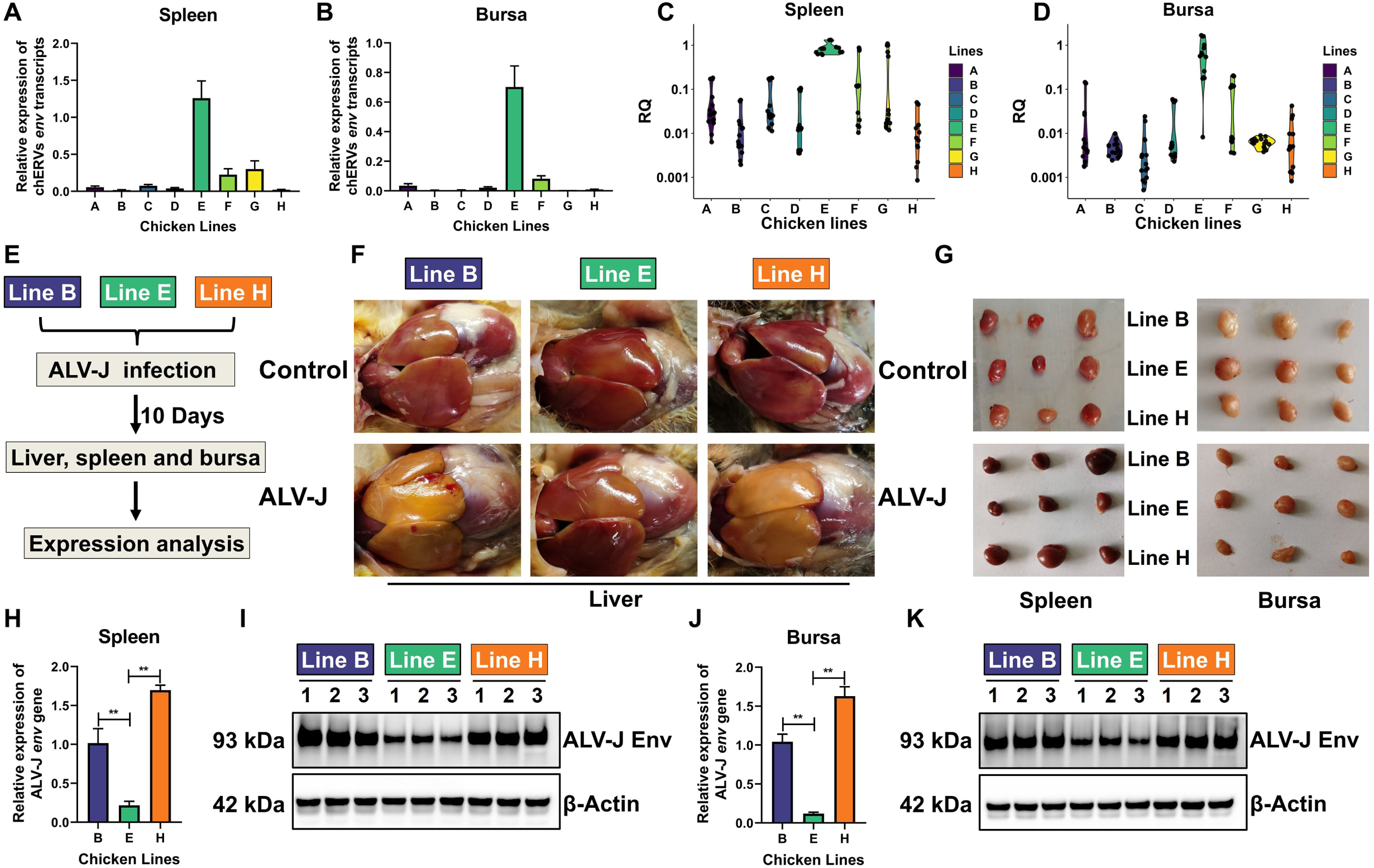
Endogenous retroviruses exert genetic resistance to the highly homologous exogenous ALV-J infection in chickens. RT-qPCR analysis of chERVs *env* transcripts expression in the central immune organs spleen (**A**) and bursa (**B**) from 8 chicken lines. Violin plot shows the distribution of the chERVs *env* transcripts expression data from 8 chicken lines spleen (**C**) and bursa (**D**). (**E**) Flow chart of ALV-J infection experiments. (**F**) The macroscopic appearance of the liver tissues from ALV-J infected chicken line B, line E and line H. (**G**) The macroscopic appearance of the spleen and bursa tissues from ALV-J infected chicken line B, line E and line H. RT-qPCR (**H**) and western blot (**I**) analysis of ALV-J *env* gene expression in ALV-J infected spleen tissues from chicken line B, line E and line H. RT-qPCR (**J**) and western blot (**K**) analysis of ALV-J *env* gene expression in ALV-J infected bursa tissues from chicken line B, line E and line H.

### chERVs inhibit ALV-J infection in macrophages

Next, we examined viral replication in macrophages, which is susceptible to ALV-J infection and associated viral immune evasion. Bone marrow-derived macrophages (BMDMs) were isolated from ALV-J-susceptible (Line B) chickens and ALV-J-resistant (Line E) chickens, and then infected ALV-J (MOI=5) for 36 h. Results showed that the expression of ALV-J *env* gene at both mRNA and protein levels in BMDMs of ALV-J-susceptible (Line B) chicken was significantly higher than that in ALV-J-resistant (Line E) chickens (Figure 2A and B). ALV-J-susceptible (Line B) chicken BMDMs were also treated with the DNA methyltransferase inhibitor 5-Aza-dC to activate the expression of chERVs-*env* transcripts and then infected ALV-J for 36 h. 5-Aza-dC treatment significantly inhibited the expression of ALV-J *env* gene at both mRNA and protein levels in ALV-J-susceptible (Line B) chicken BMDMs (Figure 2C-E). These results suggested chERVs-*env* transcripts is associated with host viral resistance in macrophages.

**Figure 2.**
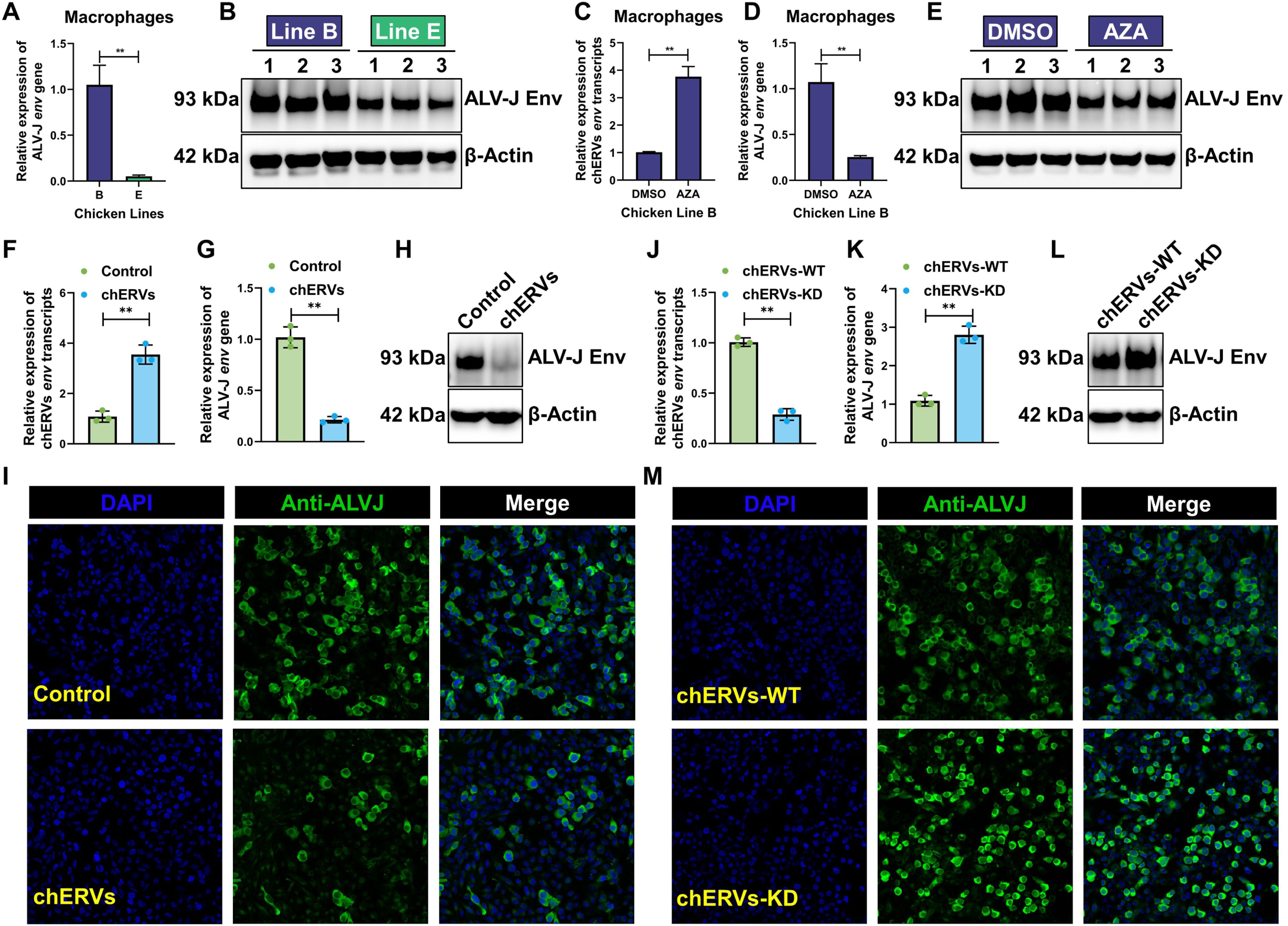
chERVs inhibit ALV-J infection in macrophages. RT-qPCR (**A**) and western blot (**B**) analysis of ALV-J *env* gene expression in ALV-J infected bone marrow-derived macrophages (BMDMs) from chicken line B and line E. RT-qPCR analysis of chERVs *env* transcripts expression in BMDMs from chicken line B with demethylation by 5-aza-2′-deoxycytidine (AZA). RT-qPCR and western blot (**E**) analysis of ALV-J *env* gene expression in ALV-J infected BMDMs from chicken line B with demethylation by AZA. (**F**) RT-qPCR analysis of chERVs *env* transcripts expression in the chicken macrophage cell line HD11 cells, transfected with chERVs-*env* or the control for 48 h. RT-qPCR (**G**) and western blot (**H**) analysis of ALV-J *env* gene expression in ALV-J infected in HD11 cells, which firstly infected with the ALV-J virus at MOI of 5 for 2 h and then transfected with chERVs-*env* or the control for another 48 h. (**I**) Confocal immunofluorescence microscopy analysis of ALV-J *env* gene expression in macrophages transfected with chERVs-*env* or the control. HD11 cells were incubated with the anti-ALV-J envelope protein (JE9 antibody) and then stained with goat anti-mouse IgG conjugated with the Alexa Fluor 488 dye (Sigma-Aldrich). The nuclei were stained with DAPI dye (Sigma-Aldrich). The pictures were captured and merged with a Leica SP8 confocal microscope (20×). (**J**) RT-qPCR analysis of chERVs *env* transcripts expression in chERVs-WT and -KD macrophages. RT-qPCR (**K)** and western blot (**L**) analysis of ALV-J *env* gene expression in chERVs-WT and -KD macrophages infected with the ALV-J virus at MOI of 5 for 48 h. (**M**) Confocal immunofluorescence microscopy analysis of ALV-J *env* gene expression in chERVs-WT and -KD macrophages. Error bars represent the s.d., n=3. *P < 0.05 and **P < 0.01 (two-tailed Student’s t-test).

To further determine the antiviral function of chERVs-*env* transcripts during ALV-J infection, we assessed the influence of chERVs-*env* transcripts on ALV-J proliferation in chicken macrophage cell lines HD11 cells. Overexpression of chERVs-*env* transcripts in macrophages significantly inhibited the expression of ALV-J *env* gene at both mRNA and protein levels at a multiplicity of infection (MOI) of 5 (Figure 2F-I). Conversely, knockdown of chERVs-*env* transcripts significantly increased the expression of ALV-J *env* gene at both mRNA and protein levels in macrophages (Figure 2J-M). Our data collectively indicate that chERVs-*env* transcripts may possess a function in antiviral defense against the oncogenic retrovirus ALV-J infection in chicken macrophages.

### chERVs knockdown block MARCO-mediated inflammatory response

To explore molecular changes in macrophages influenced by chERVs, we performed RNA-sequencing (RNA-seq) on wild-type and mutant macrophages (Figure 3A). Differential expression analysis revealed that more than 2000 genes (all of them FC>2 and q-value<0.05) were differentially expressed in chERVs-KD macrophages (Figure 3B). The top differentially expressed genes (DEGs) including GPR34, MARCO, SMOC1, TACR3, S100A9 and GPX3. We then conducted gene set enrichment analysis (GSEA) using hallmark pathways on all expressed genes in GSE270098 to identity the core genes and key pathways involved in antiviral immunity. GSEA results indicated that the top five significantly enriched hallmark pathways included interferon gamma response, inflammatory response, allograft rejection, interferon alpha response and coagulation (Figure 3C). Chord diagram further showed that the largest number of DEGs were enriched in the inflammatory response (Figure 3D). Furthermore, the top three DEGs, including TACR3, MARCO and SLC7A2, were all in the inflammatory response pathway.

**Figure 3.**
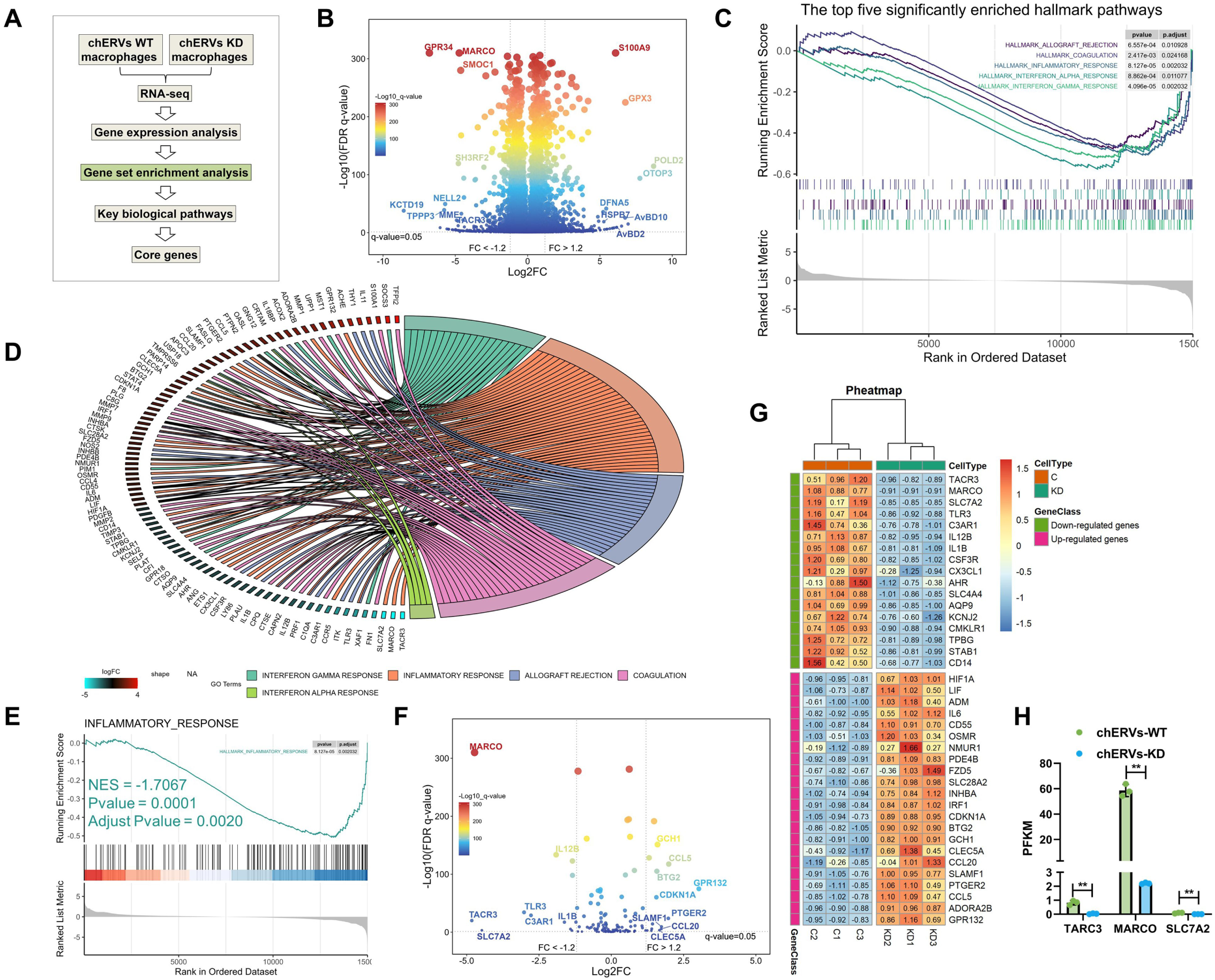
chERVs knockdown blocks the MARCO-mediated inflammatory response in macrophages. (**A**) Flowchart of analyzing core genes associated with the key biological pathways in chERVs WT and KD macrophages. (**B**) Volcano plot shows the distribution of all genes detected in chERVs WT and KD macrophages. (**C**) GSEA analysis results of the top five significantly enriched hallmark pathways between chERVs WT and KD macrophages dataset. (**D**) Chord diagram showing the relationship between the top five significantly enriched hallmark pathways and differentially expressed genes. The colored chords linking each gene to the certain hallmark pathways it is related to, and red-to-cyan coding next to the genes indicates logFC. The red and cyan colours represent the up- and down-regulated expression, respectively. (**E**) GSEA analysis results of the hallmark inflammatory response between the chERVs WT and KD macrophages dataset. (**F**) Volcano plot shows the distribution of the hallmark inflammatory response genes detected in chERVs WT and KD macrophages. (**G**) Pheatmap analysis shows the DEGs of the hallmark inflammatory response detected in chERVs WT and KD macrophages. (**H**) Gene expression analysis of TACR3, MARCO and SLC7A2 in chERVs WT and KD macrophages.

Next, we focused on the inflammatory response to reveal the mechanism of antiviral immunity in macrophages medicated by chERVs. GSEA analysis results showed a significant negative correlation (NES = -1.7067 and Adjust P-value = 0.0020) between chERVs knockdown and inflammatory response (Figure 3E). Volcano plot shows the distribution of the hallmark inflammatory response genes (including the top three DEGs TACR3, MARCO and SLC7A2) detected in chERVs WT and KD macrophages (Figure 3F). In the pheatmap of inflammatory response, the expression of genes associated with inflammation, including MARCO, toll-like receptor 3 (TLR3), Interleukin 1β (IL-1β) and IL-12β were significantly downregulated in chERVs-KD macrophages (Figure 3G). Amongst the top three DEGs, MARCO is the most abundant mRNA transcripts while the other two (TACR3 and SLC7A2) were extremely low in macrophages (Figure 3H). MARCO is the class A macrophage scavenger receptor on the cell surface of macrophages and mostly associated with inflammatory response. Thus, we speculated that MARCO might be a major player for inflammatory response regulated by chERVs.

### MARCO knockout block inflammatory response

To further investigate the function of MARCO in the inflammatory response, we knocked out the MARCO gene in HD11 cells using the CRISPR/Cas9 technique (Figure 4A). The sequencing results showed that a reading frame shift mutant with a 7 bp deletion (TTTCAGA) in exon 1 of chicken MARCO gene was generated in the sgRNA targeted HD11 cells (Figures 4A). Immunoblot analysis further confirmed the complete loss of MARCO protein in the MARCO-KO HD11 cells (Figure 4B). RNA-seq was then performed on MARCO-KO macrophages to explore molecular mechanisms of MARCO. Differential expression analysis revealed that more than 800 genes (all of them FC>2 and q-value<0.05) were differentially expressed in MARCO-KO macrophages (Figure 4C). The top DEGs including MARCO, AKR1E2, COBLL1, PREX2, CACNA1D, MAMDC2, ABCA9, PCBP2, ADAM12, FGFR3 and MYO7B.

**Figure 4.**
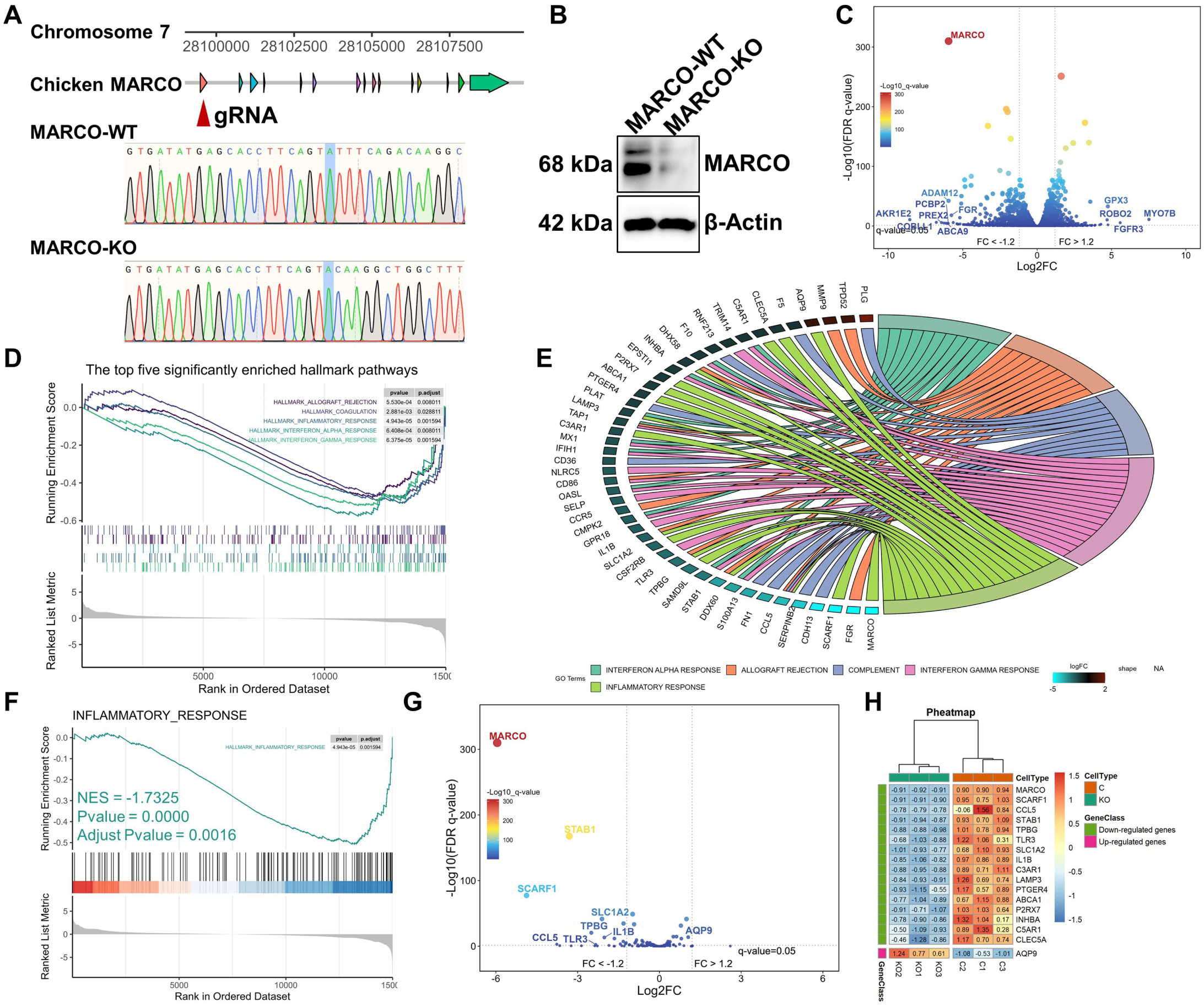
MARCO knockout blocks the inflammatory response pathway in macrophages. (**A**) Knocked out the MARCO gene in the chicken macrophage cell line HD11 cells using the CRISPR/Cas9 technique. (**B**) Western blot analysis of MARCO gene expression in MARCO WT and KO macrophages. (**C**) Volcano plot shows the distribution of all genes detected in MARCO WT and KO macrophages. (**D**) GSEA analysis results of the top five significantly enriched hallmark pathways between the MARCO WT and KO macrophages dataset. (**E**) Chord diagram showing the relationship between the top five significantly enriched hallmark pathways and differentially expressed genes. The colored chords linking each gene to the certain hallmark pathways it is related to, and red-to-cyan coding next to the genes indicates logFC. The red and cyan colours represent the up- and down-regulated expression, respectively. (**F**) GSEA analysis results of the hallmark inflammatory response between the MARCO WT and KO macrophages dataset. (**G**) Volcano plot shows the distribution of the hallmark inflammatory response genes detected in MARCO WT and KO macrophages. (**H**) Pheatmap analysis shows the DEGs of the hallmark inflammatory response detected in MARCO WT and KO macrophages.

GSEA results on all expressed genes in GSE264029 indicated that the top five significantly enriched hallmark pathways included allograft rejection, coagulation, inflammatory response, interferon alpha response and interferon gamma response (Figure 4D). Chord diagram further showed that the largest number of DEGs, including MARCO, SCARF1, CCL5, STAB1, TPBG, TLR3 and IL-1β, were enriched in the inflammatory response (Figure 4E). Furthermore, there was a significant negative correlation (NES=-1.7325 and Adjust P-value = 0.0016) between MARCO knockout and inflammatory response (Figure 4F). Volcano plot shows the distribution of the hallmark inflammatory response genes detected in MARCO WT and KO macrophages (Figure 4G). Pheatmap analysis shows 17 DEGs of the hallmark inflammatory response, and most of them were significantly downregulated in MARCO-KO macrophages (Figure 4H). These results suggested that MARCO knockout weaken the inflammatory response of macrophages.

### chERVs regulate MARCO-TLR3-IL-1β inflammatory response

Next, we aimed to identify key genes involved in inflammatory response that relay the anti-viral effect of chERVs-*env* transcripts on macrophages. We did pairwise comparisons between samples (WT vs. chERVs; WT vs. MARCO) to looked for overlap genes involved in the inflammatory response between these datasets. This revealed 6 common differentially expressed genes, including MARCO, STAB1, TLR3, C3AR1, IL-1β and TPBG (Figures 5A). To better understand the regulatory mechanisms employed by chERVs, the protein– protein interactions (PPI) network(s) of 6 common DEGs was constructed through STRING Interactome database. The network contained 6 nodes, 3 edges and 1 seed, and the 4 core genes (MARCO, TLR3, C3AR1, IL-1β) were clustered in the network (Figure 5B). Among these candidate genes, C3AR1 are important components of the complement system whereas MARCO is a class A scavenger receptor on the cell surface of macrophages. MARCO mediate double-stranded RNA (dsRNA) uptake through binding to viral dsRNA at the cell surface and to enable this dsRNA to interact with the important dsRNA recognition receptor TLR3 in the endosome (Carpentier *et al*, 2019; MacLeod *et al*, 2013). Activation of TLR3 induces inflammatory response (such as increases IL-1β expression) and interferon response, and has been frequently linked with the antiviral immune response. Therefore, it is highly likely that MARCO exerts an inflammatory response through TLR3 and IL-1β.

**Figure 5.**
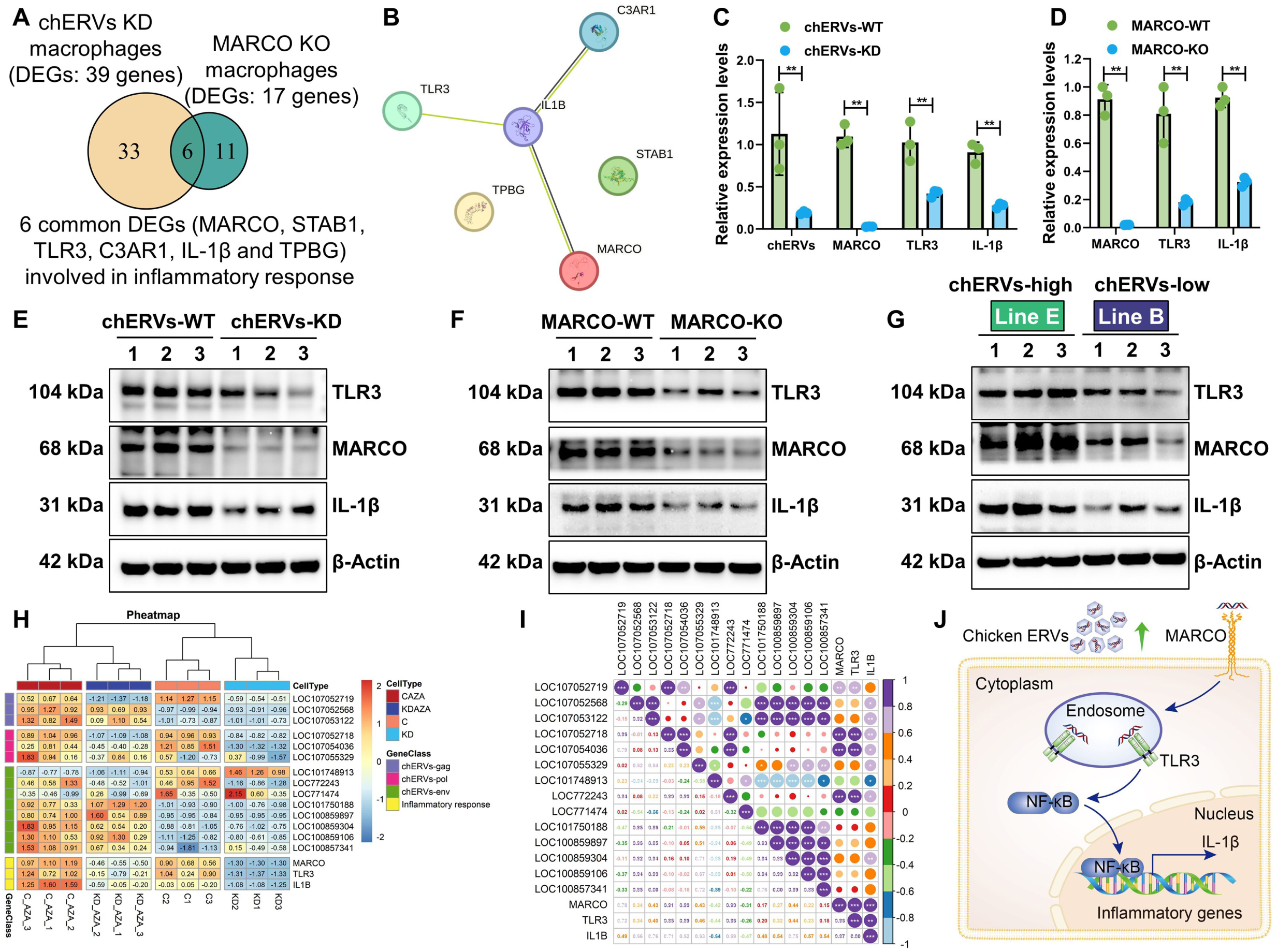
chERVs regulate the MARCO-TLR3-IL-1β inflammatory response pathway in macrophages. (**A**) Transcriptome analysis identifying common genes involved in the inflammatory response from DEGs in the chERVs KD or MARCO KO macrophages identified by RNA-seq. (**B**) The PPI network of 6 common DEGs was constructed through STRING Interactome database. (**C**) RT-qPCR analysis of chERVs-*env*, MARCO, TLR3 and IL-1β gene expression in the chERVs-WT or -KD macrophages. (**D**) RT-qPCR analysis of MARCO, TLR3 and IL-1β gene expression in the MARCO-WT or -KO macrophages. (**E**) Western blot analysis of MARCO, TLR3 and IL-1β gene expression in the chERVs-WT or -KD macrophages. (**F**) Western blot analysis of MARCO, TLR3 and IL-1β gene expression in the MARCO-WT or -KO macrophages. (**G**) Western blot analysis of MARCO, TLR3 and IL-1β gene expression in the chERVs-high (line E) or chERVs-low (line B) chicken macrophages. Pheatmap (**H**) and correlation (**I**) analysis showed that the expression of chERVs transcripts, MARCO, TLR3 and IL-1β in macrophages with chERVs knowndown or activation. (**J**) Model summarizing how chERVs regulate the MARCO-TLR3-IL-1β inflammatory response.

RT-qPCR analysis showed that the expression levels of MARCO, TLR3 and IL-1β were significantly reduced in chERVs-KD or MARCO-KO macrophages (Figures 5C and D). Western blot analysis further confirmed that MARCO deficiency significantly abolished the expression of TLR3 and IL-1β genes at protein levels in macrophages (Figures 5E and F). Furthermore, the protein expression levels of MARCO, TLR3 and IL-1β in the chERVs-high (Line E) chicken macrophages was abnormally higher than that of chERVs-low (Line B) chicken (Figures 5G). Comprehensive pheatmap and correlation analysis from all expressed genes in GSE270098 also showed that the expression of chERVs under different circumstances (activation by demethylation treatment or inhibition by CRISPR/Cas9-mediated knowndown) in the macrophages was significantly positively correlated with the expression of MARCO, TLR3 and IL-1β (Figures 5H and I). Obviously, it can be inferred from the previously findings and our results that MARCO-mediated TLR3-IL-1β inflammatory response relay the anti-viral effect of chERVs on macrophages (Figure 5J).

### ALV-J restrain MARCO-mediated inflammatory response

We next investigated MARCO-mediated TLR3-IL-1β inflammatory response patterns of macrophages infected with ALV-J. Results showed that mRNA expression of chicken MARCO was significantly and sustainably reduced in infected macrophages in a time- and dose-dependent manner (Figure 6A-D). MARCO protein expression is also dramatically declined accompanied by an increase in viral replication from 24 hours to 48 hours in infected macrophages (Figure 6E). We also observed the inhibition of TLR3 and IL-1β protein expression by ALV-J infection in infected macrophages at 48 hours (Figure 6E). ALV-J also significantly inhibited the protein expression of the MARCO, TLR3 and IL-1β in a dose-dependent manner after 48 hours post-infection (Figure 6F). These results suggest that ALV-J infection interfere with MARCO-mediated TLR3-IL-1β inflammatory response in macrophages.

**Figure 6.**
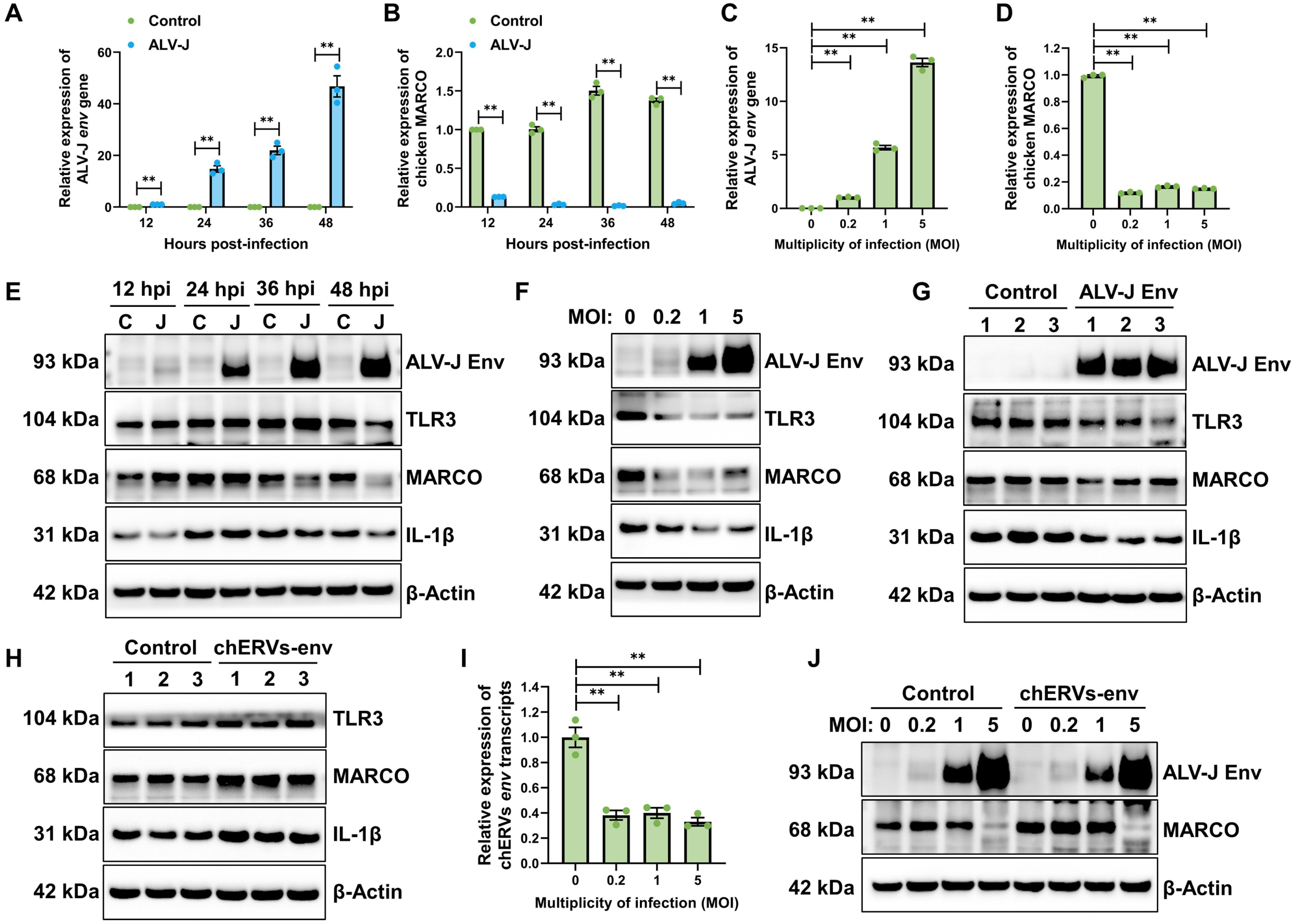
MARCO-TLR3-IL-1β pathway involved in antiviral response. RT-qPCR analysis of ALV-J *env* (**A**) and chicken MARCO (**B**) gene expression in ALV-J infected HD11 cells at 12, 24, 36 and 48 hpi. RT-qPCR analysis of ALV-J *env* (**C**) and chicken MARCO (**D**) gene expression in ALV-J infected HD11 cells infected with the ALV-J virus at MOI of 0.2, 1 and 5 for 48 h. (**E**) Western blotting analysis of MARCO, TLR3, IL-1β and ALV-J *env* protein expression in HD11 cells at 12, 24, 36 and 48 hpi. (**F**) Western blotting analysis of MARCO, TLR3, IL-1β and ALV-J *env* protein expression in HD11 cells infected with the ALV-J virus at MOI of 0.2, 1 and 5 for 48 h. (**G**) Western blotting analysis of MARCO, TLR3, IL-1β and ALV-J *env* protein expression in HD11 cells, which transfected with ALV-J *env* gene or the control for 48 h. (**H**) Western blotting analysis of MARCO, TLR3, IL-1β and ALV-J *env* protein expression in HD11 cells, with a lentiviral expression vector containing chERVs-*env* gene or EGFP. (**I**) RT-qPCR analysis of chERVs *env* transcripts expression in ALV-J infected HD11 cells infected with the ALV-J virus at MOI of 0.2, 1 and 5 for 48 h. (**J**) Western blotting analysis of MARCO and ALV-J *env* protein expression in HD11 cells with a lentiviral expression vector containing chERVs-*env* gene or EGFP. Cells were infected with the ALV-J virus at MOI of 0.2, 1 and 5 for 48 h. Error bars represent the s.d., n=3. *P < 0.05 and **P < 0.01 (two-tailed Student’s t-test).

In particular, the expression trend of MARCO is completely opposite to the viral replication level in ALV-J infected macrophages. Next, we confirm that ALV-J-encoded envelope protein is responsible for the inhibition of MARCO, TLR3 and IL-1β (Figure 6G). In contrast, chERVs-*env* overexpression activates the expression of MARCO, TLR3 and IL-1β in macrophages (Figure 6H). Furthermore, the expression levels of chERVs-*env* transcripts was significantly reduced by ALV-J infection in a dose-dependent manner, which suggested that chERVs involved in the antiviral response to ALV-J infection (Figure 6I). We also observed that chERVs promoted MARCO expression and inhibited ALV-J replication in macrophages with a lentiviral expression vector containing chERVs-*env* gene (Figure 6J). However, the protein expression of MARCO was drastically reduced in macrophages infected with higher doses (MOI=5) of ALV-J compared to macrophages infected with lower doses (MOI=0.2) or uninfected macrophages (Figure 6J). These findings suggest that ALV-J infection may disrupt the antiviral immune function of macrophages mainly through inhibiting MARCO expression and followed by TLR3-IL-1β inflammatory response.

### MARCO restrain ALV-J infection in macrophages

Persistent inhibition of MARCO after viral infection provide important insights into the molecular mechanisms underlying the interaction between ALV-J and MARCO, and may have implications for the development of therapeutic strategies for ALV-J infection. Next, we mainly investigate the antiviral function of MARCO in macrophages. It was found that overexpression of MARCO in macrophages significantly inhibited the expression of ALV-J *env* gene at both mRNA and protein levels at a MOI of 5 (Figure 7A and B). Confocal immunofluorescence microscopy analysis further confirmed that ALV-J proliferation was inhibited by MARCO overexpression in macrophages (Figure 7C). Conversely, MARCO knownout significantly increased the expression of ALV-J *env* gene at both mRNA and protein levels in MARCO-KO macrophages at a MOI of 5 (Figure 7D and E). Confocal immunofluorescence microscopy analysis further confirmed that lack of MARCO can promote the replication of ALV-J in macrophages (Figure 7F).

**Figure 7.**
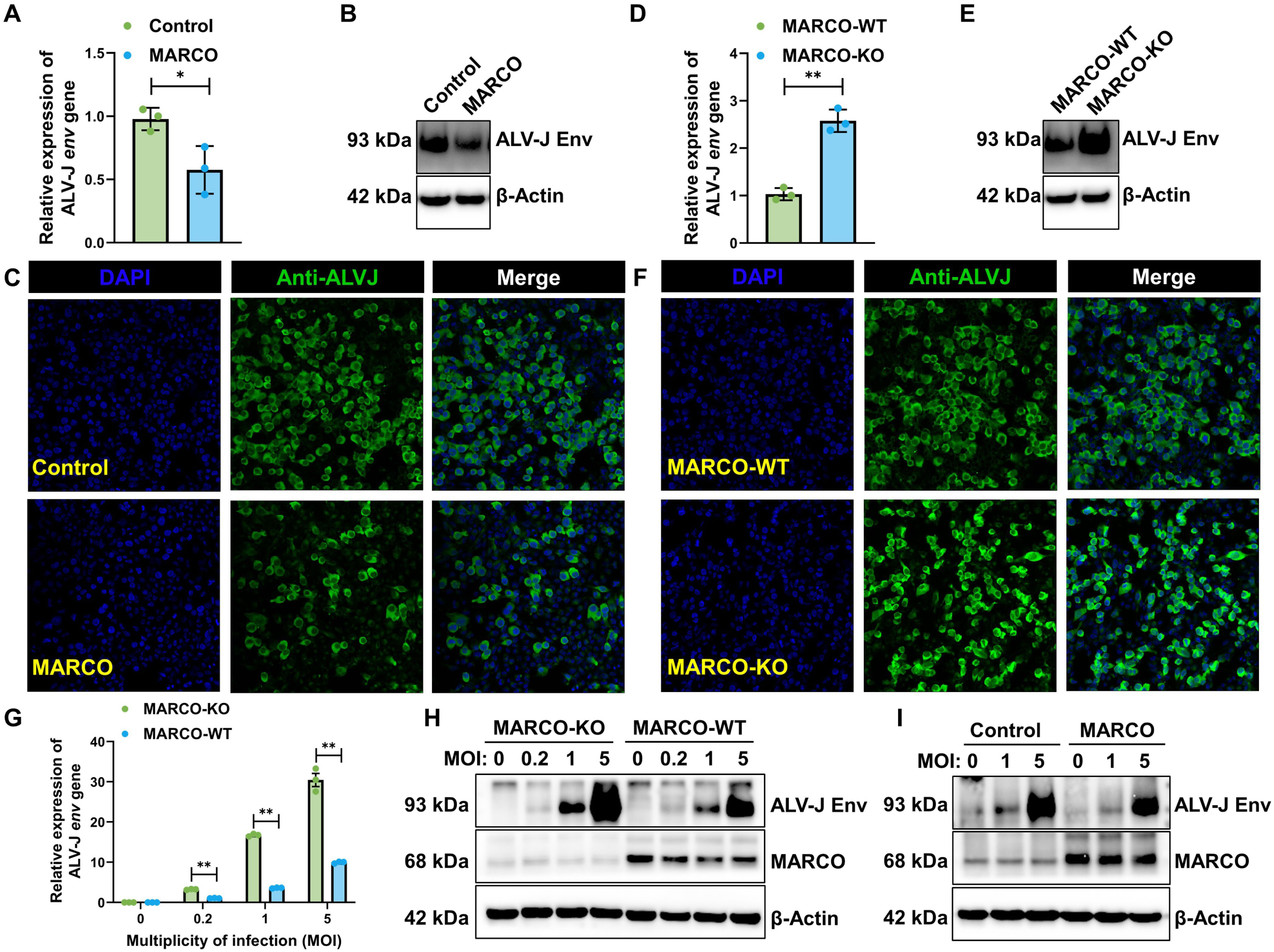
MARCO inhibit ALV-J infection in macrophages. RT-qPCR (**A**) and western blot (**B**) analysis of ALV-J *env* gene expression in ALV-J infected HD11 cells, which firstly infected with the ALV-J virus at MOI of 5 for 2 h and then transfected with MARCO or the control for another 48 h. (**C**) Confocal immunofluorescence microscopy analysis of ALV-J *env* gene expression in macrophages transfected with MARCO or the control. RT-qPCR (**D**) and western blot (**E**) analysis of ALV-J *env* gene expression in ALV-J infected MARCO-WT or -KO macrophages. (**F**) Confocal immunofluorescence microscopy analysis of ALV-J *env* gene expression in ALV-J infected MARCO-WT or -KO macrophages. (**G**) RT-qPCR analysis of ALV-J *env* gene expression in the MARCO-WT or -KO macrophages, which infected with the ALV-J virus at MOI of 0.2, 1 and 5 for 48 h. (**H**) Western blot analysis of MARCO and ALV-J *env* gene expression in the MARCO-WT or -KO macrophages, which infected with the ALV-J virus at MOI of 0.2, 1 and 5 for 48 h. (**I**) Western blot analysis of MARCO and ALV-J *env* gene expression in ALV-J infected MARCO-KO macrophages, which firstly infected with the ALV-J virus at MOI of 1 and 5 for 2 h and then transfected with MARCO or the control for another 48 h. Error bars represent the s.d., n=3. *P < 0.05 and **P < 0.01 (two-tailed Student’s t-test).

More importantly, there was a significant increased the expression of ALV-J *env* gene at both mRNA and protein levels in MARCO-KO macrophages at a MOI of 0.2, 1 and 5 (Figure 7G and H). To determine if restoring MARCO expression is sufficient to overcome ALV-J infection, we firstly infected ALV-J and then transfected with MARCO or the control in MARCO-KO macrophages. Overexpression of MARCO effectively reduced the expression of ALV-J env gene at both mRNA and protein levels in MARCO-KO macrophages at a MOI of 1 and 5 (Figure 7I), suggesting that restoration of MARCO expression is sufficient to inhibit ALV-J infection in macrophages. Our data collectively indicate that MARCO possess an important function in antiviral defense against the oncogenic retrovirus ALV-J infection in chicken macrophages.

### MARCO-mediated ligand delivery enhances TLR3-IL-1β antiviral response

To further investigate the effect of MARCO deletion on the antiviral response of macrophages, we utilized poly(I), a synthetic polyanionic ligand for TLR3 and MARCO, TLR3 ligand poly(I:C) as well as the class A scavenger receptors ligand dextran sulphate (Dxs). MARCO-WT or MARCO-KO macrophages were infected ALV-J for 24 hours, followed by stimulation with poly (I) or poly(I:C) at the concentrations of 1 and 5 μg/mL as well as Dxs at the concentrations of 1 and 5 mM for 12 hours and then detected the viral replication levels (Figure 8A). The results showed that poly (I) or poly(I:C) stimulation dose-dependently inhibited ALV-J replication in MARCO-WT macrophages (Figure 8B). The inhibition was most significant at higher doses (5 μg/mL) of poly (I) or poly(I:C), with a drastic reduction in the protein expression of ALV-J *env* gene compared to the control group. However, the ability of poly (I) or poly(I:C) to inhibit ALV-J replication was weakened in MARCO-KO macrophages. Even at high concentrations (5 μg/mL) of poly (I) stimulation of MARCO-KO macrophages, ALV-J was still able to replicate to a significant extent (Figure 8C).

**Figure 8.**
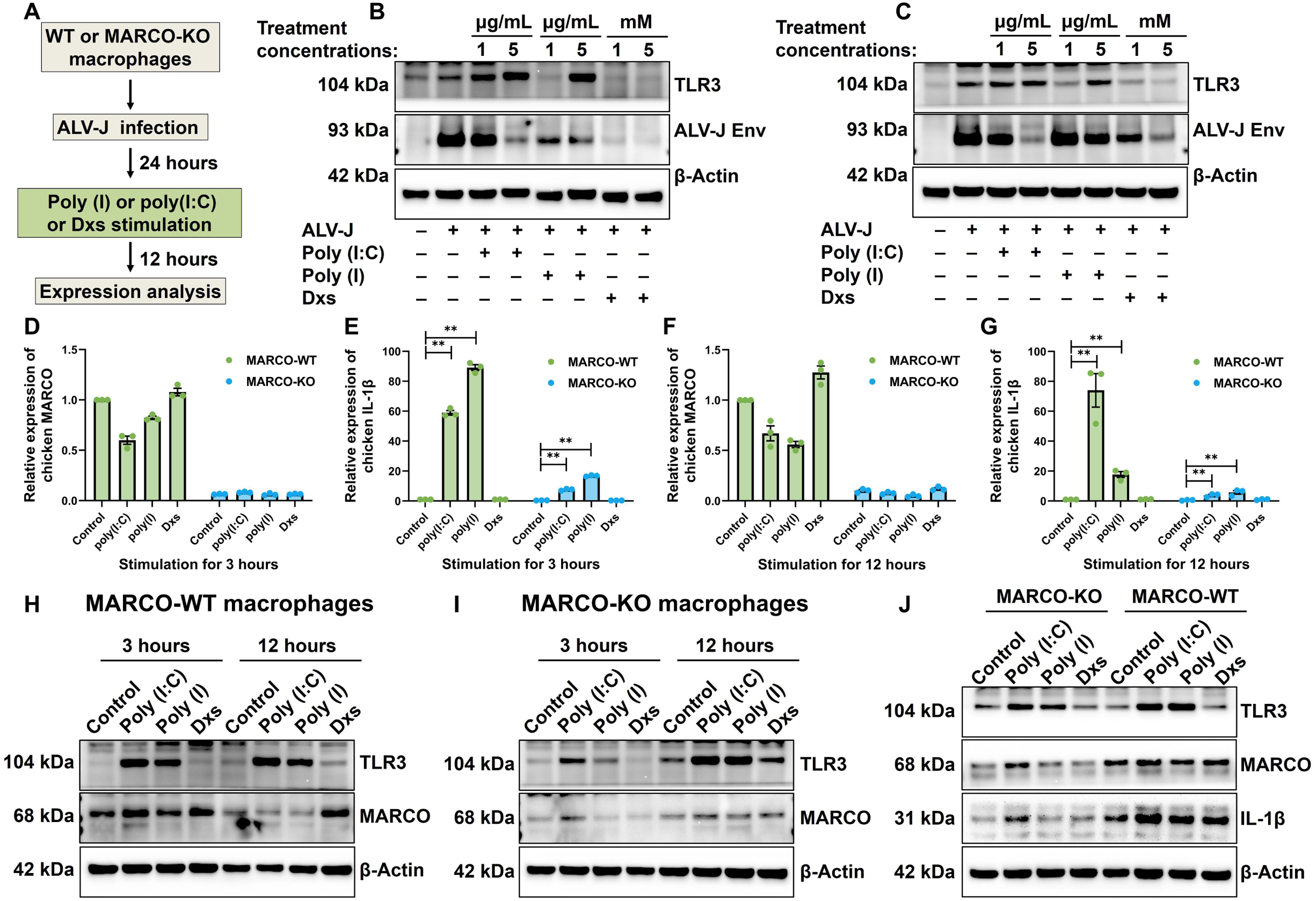
MARCO knockout block antiviral inflammatory response in macrophages. (**A**) Flowchart of analyzing the influence of drug treatment on ALV-J replication in MARCO-WT and KO macrophages. Western blot analysis of ALV-J *env* gene expression in the MARCO-WT (**B**) and MARCO-KO (**C**) macrophages, which infected with ALV-J for 24 h and then treated with poly (I) or poly(I:C) or Dxs for 12 h. RT-qPCR analysis of chicken MARCO gene (**D**) and IL-1β (**E**) expression in MARCO-WT and MARCO-KO macrophages treated with poly (I) or poly(I:C) or Dxs for 3 h. RT-qPCR analysis of chicken MARCO (**F**) and IL-1β (**G**) gene expression in MARCO-WT and MARCO-KO macrophages treated with poly (I) or poly(I:C) or Dxs for 12 h. Western blot analysis of TLR3 protein expression in MARCO-WT (**H**) and MARCO-KO (**I**) macrophages treated with poly (I) or poly(I:C) or Dxs for 3 and 12 h. (**J**) Western blot analysis of IL-1β protein expression in MARCO-WT and MARCO-KO macrophages treated with poly (I) or poly(I:C) or Dxs for 12 h. Error bars represent the s.d., n=3. *P < 0.05 and **P < 0.01 (two-tailed Student’s t-test).

Next, we investigated how the absence of MARCO in macrophages affects TLR3-mediated inflammatory response. MARCO-WT and MARCO-KO HD11 macrophages were stimulated with poly (I) or poly(I:C) or Dxs for 3∼12 hours, and the mRNA expression of IL-1β gene was measured. Results showed that poly (I) or poly(I:C) treatment for 3 or 12 hours sharply induced the expression of IL-1β in MARCO-WT HD11 cells (Figure 8D-G). Nevertheless, this induction was greatly constrained in the poly (I) or poly(I:C) stimulated MARCO-KO macrophages (Figure 8D-G). In MARCO-WT HD11 cells, poly (I) or poly(I:C) treatment for 3 or 12 hours excessively enhanced TLR3 expression at protein levels, but abnormal expression of TLR3 was only observed in MARCO-KO macrophages at 12 hours (Figure 8H and I). These results suggested that the absence of MARCO results in a delay in TLR3 activation response to poly (I) or poly(I:C) stimulation. Consistent with this, IL-1β production was abolished in the poly (I) or poly(I:C) treated MARCO-KO macrophages compared with MARCO-WT macrophages, which indicating that MARCO is required for efficient IL-1β production (Figure 8J). Endosomal TLR3 recognizes exogenous dsRNA with internalization through ligand delivery, while MARCO mediates dsRNA uptake at the cell surface. Therefore, the above findings demonstrated that MARCO-mediated ligand delivery enhances TLR3-mediated pro-inflammatory cytokine IL-1β production and antiviral response.

### MARCO-mediated IL-1β expression is responsible for viral inhibition

We have concluded that MARCO plays an important role in pro-inflammatory cytokine IL-1β production. To confirm the effect of MARCO in regulating IL-1β expression, we utilized poly (I) or poly(I:C) to stimulate macrophages which the absence of MARCO or chERVs downregulation (which results in MARCO downregulation). As expected, the mRNA expression of IL-1β was dose-dependently induced in MARCO-WT HD11 cells stimulated with poly (I) or poly(I:C) for 12 hours. Nevertheless, this induction was almost completely abolished in the poly (I) stimulated MARCO-KO macrophages and poly(I:C) stimulated chERVs-KD macrophages (Figure 9A-D). IL-1β expression was significantly attenuated in poly(I:C) stimulated MARCO-KO macrophages and poly(I) stimulated chERVs-KD macrophages. These findings suggested that MARCO absence or chERVs downregulation (which results in MARCO downregulation) impeded poly (I) or poly(I:C)-induced IL-1β production.

**Figure 9.**
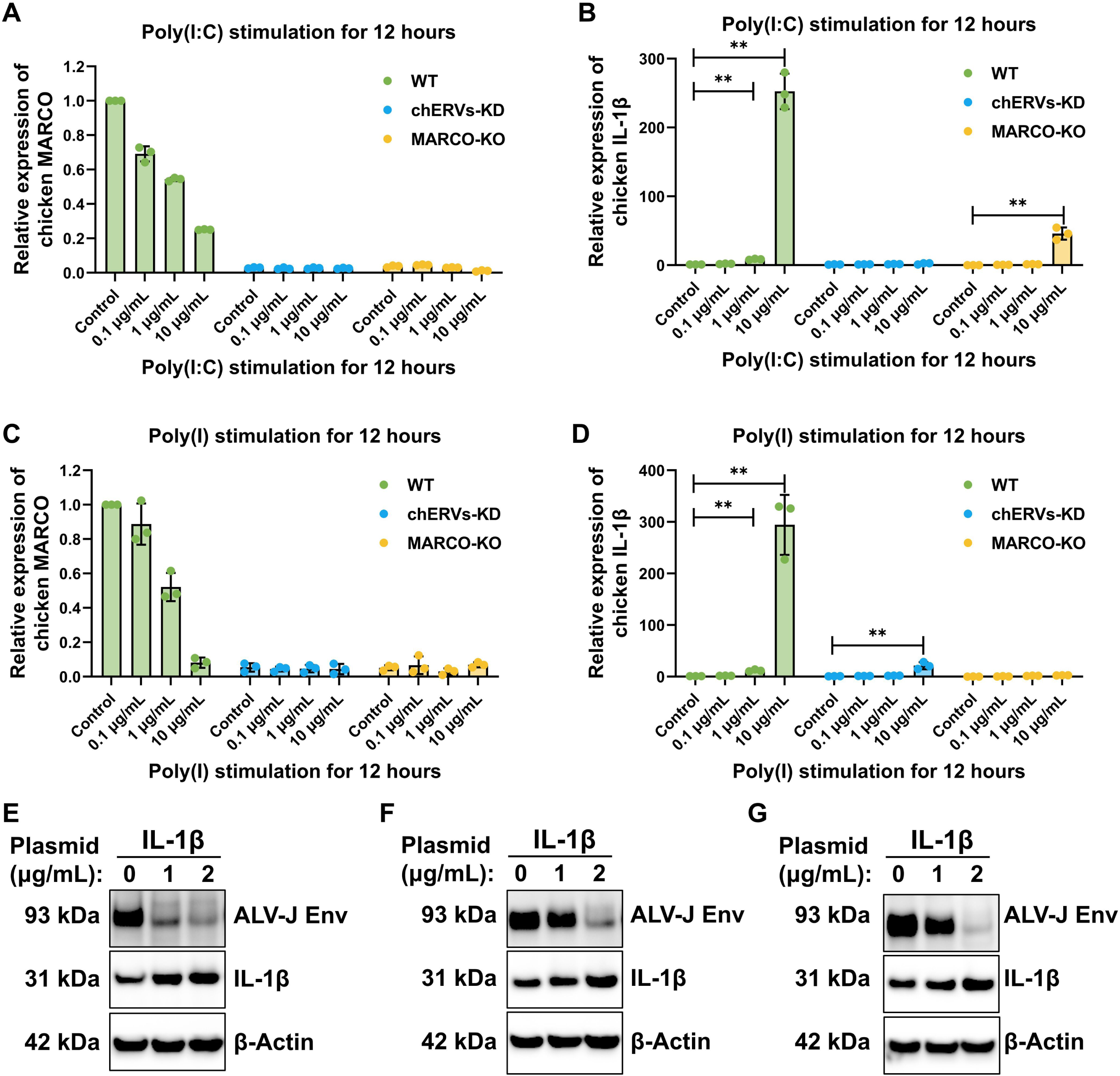
IL-1β block ALV-J replication in macrophages. RT-qPCR analysis of chicken MARCO (**A**) and IL-1β (**B**) gene expression in WT or MARCO-KO or chERVs-KD macrophages stimulated with poly(I:C) for 12 hours. RT-qPCR analysis of chicken MARCO (**C**) and IL-1β (**D**) gene expression in WT or MARCO-KO or chERVs-KD macrophages stimulated with poly (I) for 12 hours. Western blot analysis of ALV-J *env* gene expression in WT (**E**) or MARCO-KO (**F**) or chERVs-KD (**G**) macrophages, which firstly infected with the ALV-J virus at MOI of 5 for 2 h and then transfected with IL-1β for another 48 h. Error bars represent the s.d., n=3. *P < 0.05 and **P < 0.01 (two-tailed Student’s t-test).

To further investigate if IL-1β overexpression is sufficient to overcome ALV-J infection, we firstly infected ALV-J and then transfected with IL-1β or the control in MARCO−/− or chERVs-KD macrophages. IL-1β overexpression effectively reduced the protein expression of ALV-J *env* gene in WT or MARCO-KO or chERVs-KD macrophages at a MOI of 5 (Figure 9E-G). These results indicated that MARCO-mediated IL-1β expression is responsible for viral inhibition.

## Discussion

Given the widespread distribution of ERVs in the mammalian and chicken genome and their potential impact on immune regulation, investigating the association between ERVs and inflammatory response is of paramount importance. By deciphering the molecular mechanisms underlying ERVs-mediated inflammatory response modulation, we can gain novel insights into the pathogenesis of exogenous retrovirus infection and potentially uncover novel therapeutic targets for viral infection and inflammatory diseases.

The present study suggested that the chicken ERVs can stimulate the macrophage scavenger receptor MARCO which further activates TLR3. This activation triggers a signalling cascade leading to the production of IL-1β, a potent mediator of the inflammatory response. Thus, ERVs in chickens serve as “internal adjuvants”, amplifying the inflammatory response against ALV-J infection (Figure 10). Especially, we successfully employed multiple strategies to block the exogenous retroviruses ALV-J replication in chicken macrophages by drug treatments or proteins overexpression based on MARCO mediated TLR3-IL-1β inflammatory response regulated by chERVs. Overall, our findings suggest that inhibition of MARCO-mediated TLR3-IL-1β inflammatory response by exogenous retroviral infection in macrophages, which may play a role in the pathogenesis of ALV-J. Further studies are needed to fully understand the mechanisms underlying this response and its implications for the development of effective treatments for ALV-J infection.

**Figure 10.**
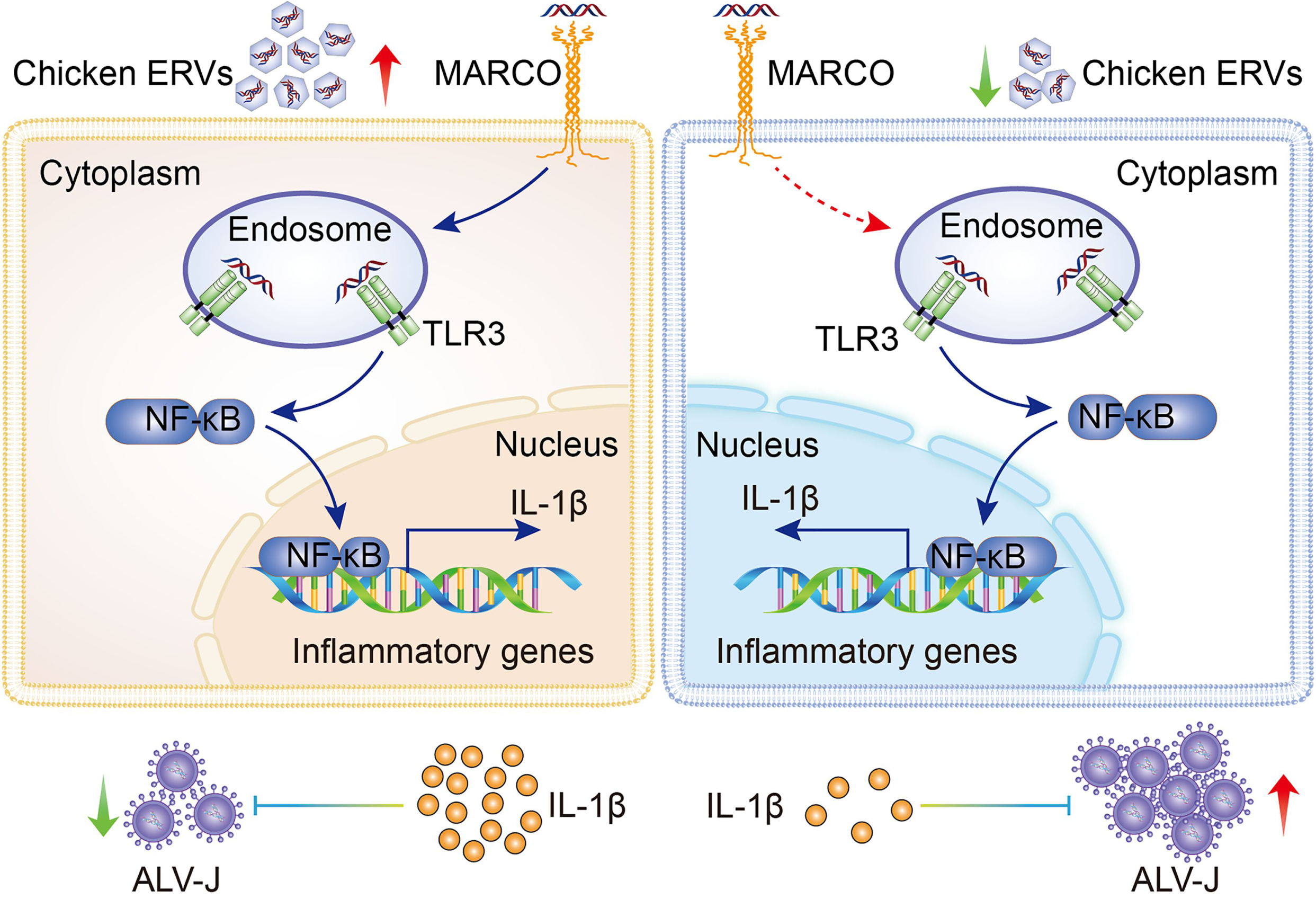
Model of chERVs antiviral inflammatory response regulation. In macrophages, high chERVs enhances the expression of MARCO and TLR3, and ultimately promoted IL-1β production, which inhibits ALV-J infection. Conversely, low chERVs attenuated the antiviral inflammatory response through reducing MARCO expression and IL-1β production.

Recently, emerging evidence has linked ERVs-derived non-coding RNAs to the innate immune system, highlighting their role in modulating the anti-viral functions of macrophages (Luo *et al*, 2022; Zhou *et al*, 2019). There are at least three proposed mechanisms for ERVs endowed the antiviral properties of macrophages through activation of innate immunity. The first is activation of interferon response via the cytosolic RNA sensors RIG-I and MDA5 (Chiappinelli *et al*, 2015; Licht, 2015; Roulois *et al*, 2015). The second is activation of interferon response via the cytosolic DNA sensors cGAS-STING (Lima-Junior *et al*, 2021) or AIM2. Here, we propose the third mechanism by which ERVs trigger inflammatory response through the scavenger receptors MARCO, the innate activation marker of macrophages.

These findings could elucidate the comprehensive mechanisms of how ERVs-derived dsRNAs are recognized by intracellular and extracellular sensors. It is well known that ERVs-derived dsRNAs can be sensed by TLR3 on the endosome or RIG-I and MAD5 in the cytoplasm and then trigger the interferon response (Chen *et al*, 2019; Chiappinelli *et al*, 2015; Licht, 2015; Luo *et al*, 2022; Roulois *et al*, 2015). However, the recognition and immune response to extracellular ERVs-derived dsRNAs are currently unclear. In this study, our results showed that ERVs-derived dsRNAs could be sensed by macrophage surface receptor MARCO and induce inflammatory response. MARCO mediate dsRNAs uptake through binding to viral dsRNAs at the cell surface and to enable these dsRNAs to interact with the important dsRNAs recognition receptor TLR3 in the endosome (Carpentier *et al*, 2019; MacLeod *et al*, 2013).

Our findings have elucidated that MARCO plays a crucial role in modulating the TLR3-IL-1β antiviral inflammatory response of macrophages. Furthermore, there was a dose- and time-dependent inhibition of the protein expression of the MARCO by ALV-J infection in macrophages. Downregulated expression of MARCO inhibited by others viruses, including SARS-CoV-2 (Haslbauer *et al*, 2023) and porcine reproductive and respiratory syndrome virus (PRRSV)(Zhang *et al*, 2023). Overexpression of MARCO significantly inhibited replication of ALV-J in macrophages, while knockout of MARCO promoted viral replication. Like other viruses, including PRRSV (Zhang *et al*, 2023) and alphaviruses (Carpentier *et al*, 2019; Carpentier *et al*, 2021; Li *et al*, 2023; Lucas *et al*, 2023), MARCO was also host restriction factor for ALV-J infection. Mechanistically, MARCO exerts inhibitory effect on ALV-J infection through regulation of TLR3-IL-1β antiviral inflammatory response. Furthermore, MARCO-mediated IL-1β expression is responsible for ALV-J inhibition. The present study has demonstrated that MARCO plays a crucial role in mediating the antiviral inflammatory response, which is employed by chicken ERVs against ALV-J infection. However, the exact mechanism by which ERVs activate MARCO and its mediated antiviral inflammatory response remains to be elucidated.

Our results also suggest an interesting evolutionary driving forces of retaining ERVs in the chicken genome. The fact that these endogenous retroviral remnants can enhance the immune response against ALV-J infection hints at a potential evolutionary advantage for the host. For examples, the domestic chickens activate the chERVs piRNA defense against ALVs (Sun *et al*, 2017), while chERVs derived lncRNAs were activated by DNA methylation inhibitors in chicken embryo fibroblasts and inhibit ALV-J replication (Chen *et al*, 2019). Our studies showed that the increased expression of ERVs concomitant with improved genetic resistance in chickens. Emerging studies also suggested a correlation between ERVs and host genetic resistance in multiple species, such as mouse (Boso *et al*, 2018), cat (Miyake *et al*, 2022; Pramono *et al*, 2024) and chicken (Mo *et al*, 2022). These findings suggest that the chicken genome may have preserved ERVs to combat external pathogens like ALV-J, this hypothesis needs to further investigation. Chicken endogenous retroviruses show highly homologous to the exogenous retroviruses, especially ALVs, and thus provide unique and ideal models for studying symbiotic relationships and the interplay among the host, endogenous and exogenous retroviruses.

In conclusion, our findings elucidated the critical role of chicken endogenous retroviruses in the MARCO-mediated antiviral inflammatory response. MARCO strikingly promoted the antiviral responses of macrophages through enhancing ligand delivery to the intracellular pathogen sensors TLR3 followed by IL-1β production. The present study highlight potential of chERVs and MARCO as genetic resistance factors and drug targets for the development of immune-modulating strategies to counter ALV-J infection. However, much work is still needed to fully comprehend these complex mechanisms and their implications in avian health and disease.

## Materials & Methods

### Ethics approval

All chicken experiments were performed in strict accordance with the recommendations provided in the Guide for the Care and Use of Laboratory Animals of Yangzhou University. The protocol was approved by the Committee on the Ethics of Animal Experiments of Yangzhou University (licence number: 06R015).

### Animal Experiments

All chickens used in this study were obtained from Changzhou Animal Disease Prevention and Control Center. Eight 1-day-old specific pathogen-free (SPF) chicken Lines (A, B, C, D, E, F, G and H) spleen and bursa were used for chERVs expression analysis. For viral infection experiments,1-day-old SPF ALV-J-susceptible (Line B) or resistant (Line E) chickens were injected intraperitoneally with ALV-J (infectious dose = 1.0×10^4^ TCID_50_/mL) or normal saline, and then kept in separate units with similar environmental conditions. At 10 days after infection (dpi), 6 chickens (3 infected and 3 uninfected control group) were sacrificed to excise the specific tissues (including liver, spleen and bursa). These tissues were then immediately stored in liquid nitrogen until gene expression analysis.

### Cell culture, virus, and plasmids

The chicken macrophage cell line HD11 were cultured in Dulbecco’s modified Eagle’s medium (DMEM; Gibco) with 10% foetal bovine serum (FBS) at 41°C in 5% CO_2_ and 95% humidity. The chicken ERVs-deficient HD11 cells was established as previous described (Liu *et al*, 2017) and maintained our laboratory. Briefly, two selection markers and a bovine growth hormone (BGH) polyA signal were inserted into the first exon of chicken ERVs ALVE1 genes via CRISPR/Cas9-mediated homology-directed repair (HDR). Next, double marker selection was employed for screening clonal cell lines with successful biallelic integration of the poly (A) signal. Finally, the efficiency of gene silencing was evaluated by RT-qPCR using primers specific for chERVs-*env* (Table 1).

**Table 1.**
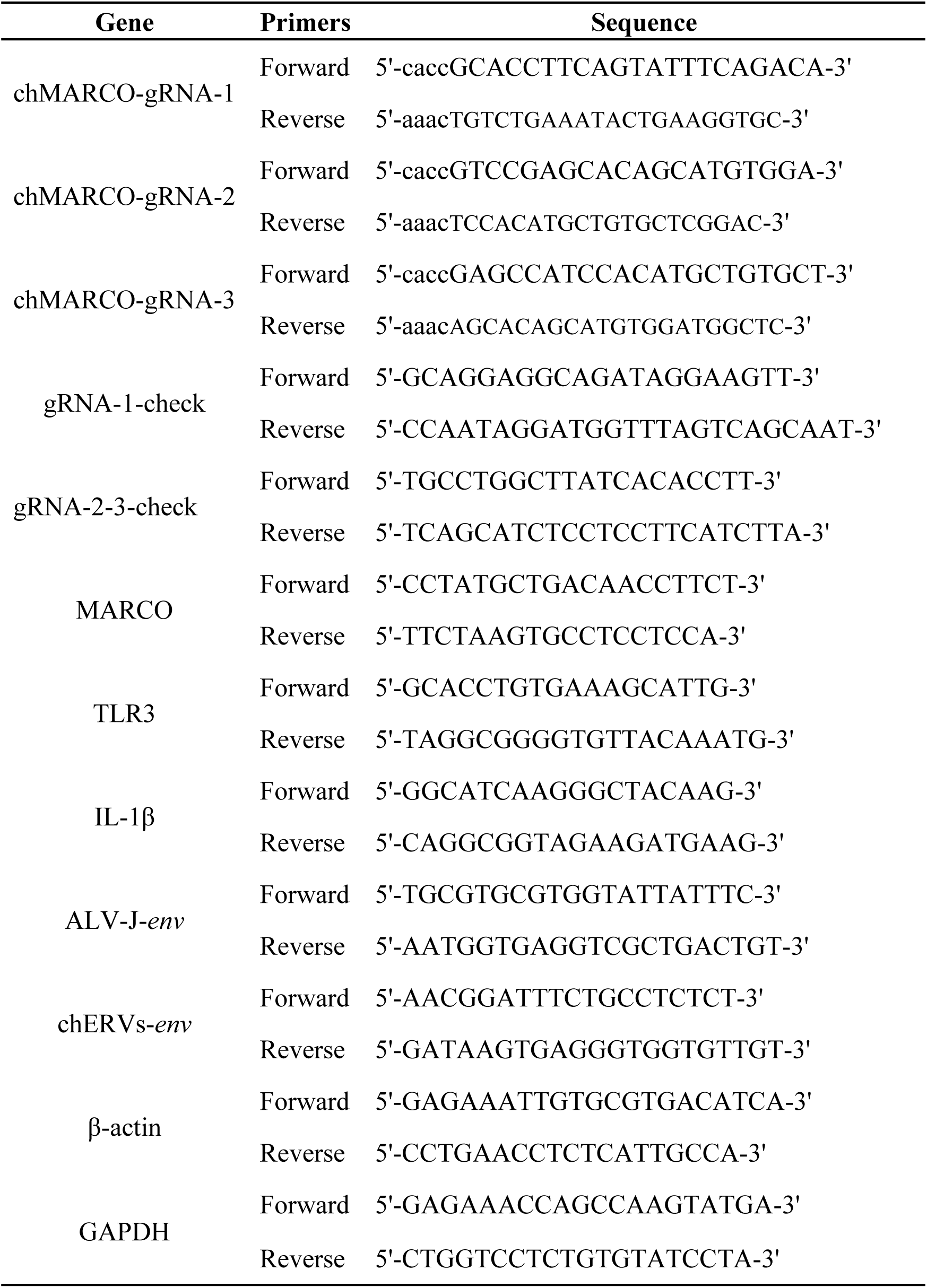
Primers used for this study.

ALV-J (the virus strain JS09GY3, GenBank accession number GU982308) was used for viral infection experiments and obtained from the Lab of Avian Preventive Medicine at Yangzhou University in China. Plasmids including the pcDNA3.1-chERVs-*env*, pcDNA3.1-MARCO, pcDNA3.1-IL-1β and pcDNA3.1-EGFP were constructed and kept in our laboratory.

### CRISPR/Cas9-mediated chicken MARCO gene knockout in HD11 cells

The MARCO-KO HD11 cell lines was established according to previous study. Briefly, Cas9-T2A-GFP plasmid and three guide RNA plasmids were firstly co-transfected into HD11 cells, respectively. After 48 h transfection, the sgRNA with higher cleavage efficiency was selected by the T7E1 assay. Cas9-T2A-GFP plasmid and the selected sgRNA were then co-transfected into HD11 cells for 48 h, and GFP-expressing cells were sorted into single wells of a 96-well dish containing DMEM with 10% FBS on a BD FACSAria™ Fusion cell sorter. Next, sorted GFP-expressing cells were cultured for 3 weeks by serial dilution to colony outgrowth. Finally, the homozygous frame-shift mutation and MARCO protein expression for these clones were identified by genomic DNA PCR (the primers used in Table 1) and western blotting to obtain knockout cell lines.

### Generating stable cell lines with lentivirus

The chERVs-*env* sequences were amplified from HD1 cDNA template with the following primers: forward 5′-TGCTCTAGACATTGGTGTGCACCTGGGT-3′ and reverse 5′-CGCGGATCCATTCCTTGCCATGCGCGAT-3′ and then cloned into the pCDH-CMV-MCS-EF1-Puro lentiviral vector. Lentiviruses containing pCDH-CMV-MCS-EF1-Puro-chERVs-*env* or EGFP were produced with a lentiviral packaging mix containing an optimized mixture of the three packaging plasmids, pLP1, pLP2, and pLP/VSVG, in 293T packaging cells. To establish stable cell lines expressing chERVs-*env* or EGFP, HD11 cells were transfected with lentivirus infection for 48 h, and stable transduced clones were generated following selection with 2 µg/mL puromycin for 1 to 2 weeks.

### Isolation of chicken bone marrow-derived monocytes (BMDMs)

21-days-old chickens were used for the generation of BMDMs. Bone marrow cells were isolated from the femur bones of the chickens. Cells were seeded in 60-mm cell culture plates and the adherent cells were collected in complete DMEM with 10% FBS for viral infection or drug treatment experiments.

### Virus infection

HD11 cells were seeded into six-well plates and infected for 3 h with the ALV-J virus at a multiplicity of infection (MOI) of 5. Upon infection, HD11 cells were washed three times with ice-cold PBS and then transfected with chERVs or MARCO or IL-1β for 48 h to analysis the virus proliferation. For the HD11 cells, ALV-J virus proliferation was analysed with Western blot, IFA and RT-qPCR at 48 h post infection.

### DNA transfection

HD11 cells were transfected with pcDNA3.1-chERVs-env, pcDNA3.1-MARCO, pcDNA3.1-IL-1β or pcDNA3.1-EGFP expression plasmids using HighGene Transfection Reagent (ABclonal, RM09014) for 48 h, and total RNA and protein were collected for gene expression analysis.

For viral infection experiments, HD11 cells were first infected with the ALV-J virus at a MOI of 5 for 3 h. Cells were then transfected with pcDNA3.1-chERVs-env, pcDNA3.1-MARCO, pcDNA3.1-IL-1β or pcDNA3.1-EGFP expression plasmids using HighGene Transfection Reagent (ABclonal, RM09014) for another 48 h and then collected for ALV-J replication analysis.

### Drugs treatment

MARCO-WT or MARCO-KO HD11 cells were treated for 3 h or 12 h with poly(I) or poly(I:C) at concentrations of 1 and 10 μg/mL or dextran sulphate (Dxs) at concentrations of 1 and 10 mM. Durgs poly(I), poly(I:C) and Dxs was purchased from Sigma-Aldrich, St. Louis, MO. Alternatively, WT or MARCO-KO or chERVs-KD HD11 cells were treated for 12 h with poly(I) or poly(I:C) at concentrations of 0.1, 1 and 10 μg/mL.

For viral infection experiments, MARCO-WT or MARCO-KO HD11 cells were infected with ALV-J (MOI=5) for 24 h, followed by treatment with poly(I) or poly(I:C) at concentrations of 1 and 10 μg/mL or Dxs at concentrations of 1 and 10 Mm for 12 h. Alternatively, WT or MARCO-KO or chERVs-KD HD11 cells were infected with ALV-J (MOI=5) for 24 h, followed by treatment with poly(I) or poly(I:C) at concentrations of 0.1, 1 and 10 μg/mL.

### RNA isolation

RNA was isolated from chicken cells or tissues using RNA-easy^TM^ Isolation Reagent (Vazyme, R701-02) according to the manufacturer’s instructions. RNA concentration was determined by Nano-300 micro spectrophotometer, and the A260/A280 ratio is used to assess RNA purity. High-quality RNA samples were used as input for RT-qPCR analysis.

### Complementary DNA synthesis and quantitative PCR

Complementary DNA was synthesized using the HiScript III RT SuperMix for qPCR (+gDNA wiper) (Vazyme, R323-01) following the manufacturer’s instructions. Quantitative PCR (qPCR) was performed using ChamQ Universal SYBR qPCR Master Mix (Vazyme, Q711-02) on a CFX Connect™ Real-Time PCR Detection System (Bio-Rad). The primers used can be found in Table 1 and Ct values were normalized to an internal control (Gapdh and β-actin).

### RNA sequencing and computational analysis

RNA sequencing libraries were prepared by BGI Genomics Co., Ltd. The reads for HD11 samples were obtained using the BGISEQ-500 or DNBSEQ-T7 sequencing platforms and filtered with SOAPnuke (v1.6.5), and then aligned to chicken reference genome (GCF_000002315.6_GRCg6a) by HISAT (v2.2.1). Differential expression analysis was performed using the DESeq2 package (v1.40.2). Gene Ontology (GO) enrichment analysis and Gene set enrichment analysis (GSEA) was performed using the clusterProfiler package in R (v.4.3.3) and RStudio (v2022.07.2+576).

### TCID_50_ Assay

DF-1 cells were seeded in growth media on each well of 96-well plates and cultured in DMEM with 5% FBS at 37 °C in 5% CO_2_ and 95% humidity. After preparing the series of dilutions at 1:10 of the original virus sample, mildly add 0.1 ml of virus dilution per well, infecting 8 wells per dilution, and then cultured for 1 week at 37°C. The viral replication in the DF1 cells was analysed with immunofluorescence assay (IFA) with mouse monoclonal antibody JE9, which is specific to the envelope protein of ALV-J at 1-week post-inoculation. The viral titer is calculated using the method of Muench and Reed.

### Western blotting

Protein was separated by 10% SDS-PAGE at 120 V for 90 min and then transferred to polyvinylidene difluoride membranes at 50 V for 150 min. Membranes were blocked in TBS-T containing 5% non-fat dry milk (BIO-RAD). Primary antibodies were incubated overnight at 4°C with agitation. The following antibodies were used to determine protein expression: rabbit polyclonal to MARCO (A10048, ABclonal), rabbit polyclonal to TLR3 (bs-1444R, Bioss), rabbit polyclonal to IL-1β (A20529, ABclonal), rabbit monoclonal to Actin (ab179467, Abcam), and mouse monoclonal antibody JE9, which is specific to the envelope protein of ALV-J. After washing extensively with TBST, secondary antibodies (anti-rabbit or anti-mouse horseradish peroxidase (HRP) conjugate, 1:10,000 dilution) were incubated for 1 h at room temperature. After washing extensively with TBST, blots were developed using enhanced chemiluminescent (ECL) detection reagents on the FluorChem Q imaging system (Protein Simple).

### Immunofluorescence staining

Cells were fixed with 4% paraformaldehyde (20 min) and permeabilized with 0.25% Triton X-100 for 15 min at room temperature. After blocking with 2% BSA for 30 min, the cells were then incubated with mouse monoclonal antibody JE9 to ALV-J overnight at 4°C, followed by a further incubation at room temperature for 45 min with an AlexaFluor®488 goat anti-mouse preadsorbed secondary (Abcam, ab150117). Nuclear DNA was labelled with DAPI dye (Sigma-Aldrich, MBD0015) at room temperature for 10 min. Images were acquired using a Leica TCS SP8 confocal microscope, and data analysis was carried out with Leica LAS AF Lite (Leica Microsystems).

### Statistical analyses

Statistical analyses were performed using GraphPad Prism version 9.51 for Windows, GraphPad Software, San Diego, California, USA, www.graphpad.com. Statistical significance was assessed using a two-tailed unpaired Student’s *t*-test with a *P* value threshold of < 0.05.

## Data Availability

The RNA-seq data in this study have been deposited in the Gene Expression Omnibus (GSE) under accession code GSE264029 and GSE270098. All data are archived at Yangzhou University and available from the corresponding author upon reasonable request.

## Author Contributions

Xuming Hu: conceptualization, formal analysis, validation, investigation, visualization, methodology, and writing—original draft. Wang Guo: validation, investigation, visualization and methodology. Huixian Wu: validation, investigation, visualization and methodology. Jinlu Liu: visualization and methodology. Xiao Han: visualization and methodology. Xujing Chen: visualization and methodology. Yu Zhang: conceptualization and formal analysis. Yang Zhang: conceptualization and formal analysis. Zhengfeng Cao: conceptualization and formal analysis. Qiang Bao: conceptualization and formal analysis. Wenxian Chai: supervision and funding acquisition. Shihao Chen: conceptualization and formal analysis. Wenming Zhao: supervision, funding acquisition, project administration. Guohong Chen: supervision, funding acquisition, project administration, and writing—review, and editing. Hengmi Cui: supervision, funding acquisition, project administration, and writing—review, and editing. Qi Xu: supervision, funding acquisition, project administration, and writing—review, and editing.

## Disclosure and competing interests statement

The authors declare no competing interests.

## Acknowledgements

This work was supported by grants from National Natural Science Foundation of China grant (32272860, 31602032), National Key Research and Development Program of China grant (2016YFC1303604), Priority Academic Program Development of Jiangsu Higher Education Institutions (Animal Science).

